# Scarcity of hosts for gregarious parasitoids indicates an increase of individual offspring fertility by reducing their own fertility

**DOI:** 10.1101/2021.03.05.434037

**Authors:** Alena Samková, Jan Raška, Jiří Hadrava, Jiří Skuhrovec

## Abstract

The gregarious parasitoid strategy allows multiple larvae to complete development in a single host due to their tolerance and/or lower mobility and thus flexibly adjust their reproductive potential amidst changing environmental conditions. Reproductive success is generally measured as the number of each mother’s offspring. We propose that with scarcity of host, for gregarious parasitoids is important the view on the fertility not only of a mother but also of her offspring (F1 generation). Due to the body size-fitness correlation, each female deliberately adjusts the clutch size, determining the offspring body size and their reproductive potential. In our study, using *Anaphes flavipes* as a model species, we showed that under a limited number of hosts, the females reduced their fertility. We propose that the lower fertility of mothers can cause higher fertility in the F2 generation using a larger offspring body while halving fertility. The females increase their individual offspring’s fertility by reducing their own fertility. Moreover, we showed that with a scarcity of hosts, the mothers increased the number of their female offspring, and thus, they obtained more offspring in the F2 generation. Additionally, other costs and benefits of the gregarious strategy in relation to superparasitism were tested and discussed.

## INTRODUCTION

Parasitoids provide a unique opportunity for studies of the evolution of reproductive behaviour^1^. In particular, gregarious parasitoids with more tolerant and/or less mobile larvae developing together in one host are suitable models^2,3^. Some combinations of the number and sex ratio of offspring developing in a single host are more advantageous than others^4,5^ and are favoured by natural selection to maximize individual fitness^6^. After finding a host, the female parasitoid decides not only on the sex (haplodiploid sex determination^7^) and the number of offspring deposited into a single host but also, in many cases of parasitoids, on their ultimate body size and fertility^8^. Because parasitoids larval development is dependent strictly on the nutrients contained in one host^9^, the amount of food obtained during larval development positively correlates with their body size^10,11,12^ and their body size determines their fertility^11,13^.

Under optimal conditions, such as a sufficient number of hosts and a sufficient number of eggs a female has available^14^, she can choose the optimal clutch size per single host without a trade-off between the clutch size and the offspring body size (Samková et al. 2019a). Even distribution of offspring among a sufficient number of hosts ensures a large offspring body size and their subsequent high fertility^8, 15^. With a scarcity of hosts, the parasitoids female faces a trade-off between the clutch size and the body size of the offspring, and she chooses a reproductive strategy of smaller offspring^15^.

In the natural environment, the parasitoids female faces intraspecific and interspecific competition for hosts^16,9,17^. In competition with predators of the host, the parasitoids can recognize predators’ chemical marks^18,19^ and, based on this recognition, avoid parasitation^17,18^ or change their clutch size^15^.

On the other hand, the competition between parasitoids is, in many cases, the result of superparasitism, repeated parasitism of the same host by the same^20^ or different parasitoid species^9^. The costs of superparasitism are a smaller body size^12^ and a reduction in body size, longevity and/or fertility of the offspring^11,13,21^. For this reason, many species of parasitoids are able to recognize an already parasitized host^22,23,20,24^ and accordingly to change their gregarious reproductive strategy^9,25,26,21^. For example, when the probability of superparasitism is high, the first female should lay a smaller number of offspring into one host, while the second female lays even fewer offspring into the same host^27,9^. The total fitness gained from one host may still increase with a higher number of parasitoids, at least until a maximum is reached^9^ but exceeding the capacity of the host causes starvation of the larvae^28^. According to a generally accepted hypothesis, avoiding superparasitism is adaptive when enough unparasitized hosts are available, and it is an adaptive strategy for individual parasitoids to compete for hosts^29,9^. However, for many cases of gregarious parasitoids, superparasitism can have a direct intergenerational impact. The smaller offspring body size in the F1 generation reduces the fertility in the F2 generation^11,30,8^.

The idiobiont gregarious parasitoid *Anaphes flavipes* (Förster, 1841) (Hymenoptera: Mymaridae), is a suitable model to study the decision-making process about intergenerational impacts of reproductive strategies with relation to the scarcity of hosts because the body size— fitness correlation applies to *A. flavipes*, i.e., the clutch size determines the offspring body size, and the body size affects their fertility^8^. We assumed possible changes in reproductive strategy (total number and sex ratio of offspring, clutch size and the total number of parasitized hosts) due to superparasitism of the food resources, because superparasitism has previously been observed for *A. flavipes*^31^. They also change their reproductive strategy under different external conditions such as the host population density and the presence of predators of host eggs^15^. Their host spectrum includes the genera *Lema* and *Oulema* (Coleoptera: Chrysomelidae), especially economically important crop pests in Europe and North America, and the cereal leaf beetle *O. melanopus* Linnaeus, 1758^31,32,33^.

In this study, we focused on the response of a gregarious reproductive strategy due to superparasitism in relation to a scarcity of hosts using *A. flavipes* as a model species. We point out the importance of the effect of changes in reproductive strategy on the hypothetical fertility in the F2 generation. We tested two different scenarios: (1) ***Different numbers of founders***. With a higher number of founders and increased pressure towards a superparasitic strategy, the clutch size increased, but fertility per each female decreased^9^. We expected similar trends for the reproductive behaviour of *A. flavipes;* (2) ***Same number of founders with a different design***. We measured the reproductive strategy of four groups with different designs of parasitation and interactions of the same number of founders. *Group 1*. We assumed reduced fertility per each female with a higher clutch size because the wasps were with the host eggs together and had visual contact with each other. Under these conditions, females assume that the host has been parasitized or will be superparasitized^14^ and reduce their own fertility^34,9^; *Group 2a*. We assumed a possible difference in wasp reproductive strategy because in this group, the first female is alone with the host eggs; she has a higher fertility and clutch size because she does not assume superparasitization of the host. However, a second female is then added to the Petri dish, and she can deliberately reduce her own fertility because she assumes a parasitized host using visual information about the first female^14^; *Group 2b*. This group is similar to the previous one, but both females are with the host eggs separately. The second female has no visual information about the first female and she can recognize the parasitized host only by chemical marks^35^. In general, there is a strong pressure to recognize parasitized hosts^36^, and the second founder lays smaller clutch^37,34^ or avoids superparasitization if there are enough unparasitized hosts^29,9^. However, females of some parasitoids species cannot recognize the parasitized host^27,38^ and lay a higher clutch size because they consider the host to be unparasitized. We expected the second *A. flavipes* female to reduce her clutch size because Anderson & Paschke^31^ showed that *A. flavipes* prefers unparasitized hosts to parasitized hosts; *Group 2c*. Here, the females were with the host eggs separately, but the timing between parasitation was long (24 hours). We expected a possible lower clutch size and fertility of wasps with a long latency between parasitation compared to wasps with a short latency, because of the survival of the offspring of the second founders is time-dependent^39^, thus the second female can reject the parasitized host after a critical time from the parasitation by the first female^26^.

## RESULTS

### (1) Different number of founders

Different numbers of founders with a constant amount of hosts (*Groups 1, 2* and *3*) affected the reproductive strategy of *A. flavipes*. The total number of offspring (LM, F1,59 = 20.537, P < 0.001, n = 61, Fig. 1a) and the total number of parasitized host eggs per Petri dish (GLM-b, χ^2^_1_ = 6.262, P = 0.012, n = 61, Fig. 1b) increased with higher numbers of founders. However, the relative fertility per female decreased (LM, F_1,59_ = 97.493, P < 0.001, n = 61, Fig. 1a - shaded part of the columns). The number of offspring developing in one host increased with a higher number of founders (LMM, χ^2^_1_ = 23.01, P < 0.001, n = 61, Fig. 1c). The sex ratio of offspring per dish was more female-biased when the number of founders was higher (GLMM-b, χ^2^_1_ = 4.869, P < 0.001, n = 61, Fig. 1d).

**Fig. 1.**
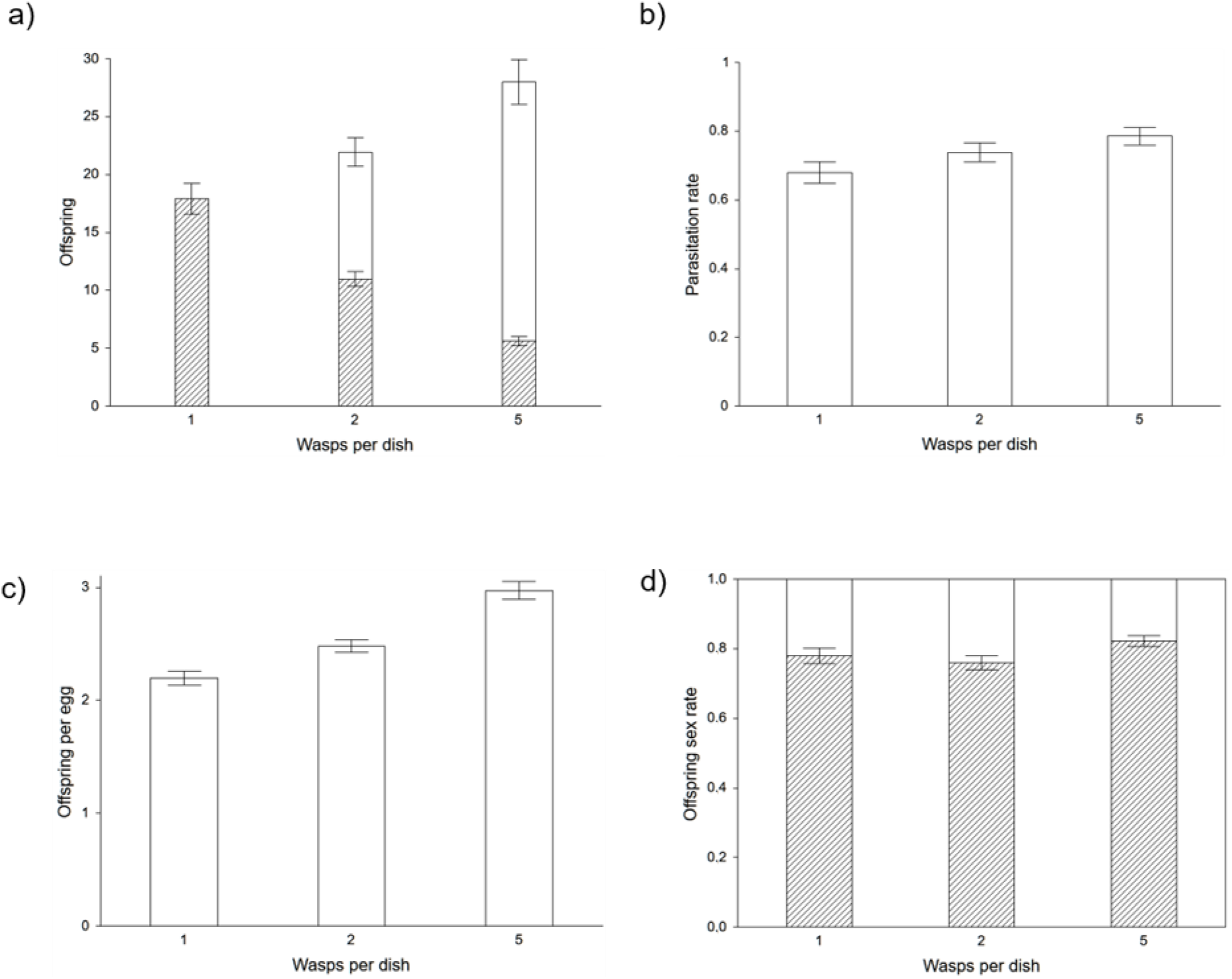
The relationship between the number of founders per dish and the total number of offspring (a) (the total number of offspring relative to one female - shaded part of the columns), the total number of parasitized host eggs (b), the offspring per one parasitized host egg (c) and the percentage of female offspring (d).

### (2) Same number of founders with a different design

We compared the four groups with the same number of founders with different designs for parasitation with each other. *Groups 2, 2a, 2b* and *2c* (Fig. 2) had a statistically significant effect only on the clutch size. The clutch size in *Group 2* was higher compared to *Group 2b* (LMM, χ^2^_1_ = 8.581, P = 0.003, n = 40, Fig. 2c) and the clutch size of *Group 2c* was lower compared to *Group 2* (Gamma-GLMM, χ^2^_1_ = 7.461, P = 0.006, n = 41, Fig. 2c) and *Group 2a* (Gamma-GLMM, χ^2^_1_ = 4.131, P = 0.042, n = 41, Fig. 2c). Differences between sex ratios, total numbers of offspring per dish or total numbers of parasitized hosts per dish among these four groups were not statistically significant (Fig. 2a, b, d). When merging the four groups according to interactions between founders (*Groups 2 + 2a* versus *2b + 2c*, Fig. 2), the total number of offspring per one Petri dish (LM, F1,79 = 4.789, P = 0.032, n = 81, Fig. 2a) and the clutch size (Gamma-GLMM, χ^2^_1_ = 9.656, P = 0.002, n = 81, Fig. 2c) were higher in Petri dishes in which the wasps could interact with each other (*Groups 2 + 2a*) than in dishes in which the wasps did not meet (*Groups 2b + 2c)*. The number of parasitized eggs per one Petri dish (LM, F1,79 = 0.013, P = 0.911, n = 81, Fig. 2b) and the sex ratio of the offspring (GLMM-b, χ^2^_1_ = 2.069, P = 0.15, n = 81, Fig. 2d) did not differ significantly among the treatments. All values are shown in the table in Supplementary material 2).

**Fig. 2.**
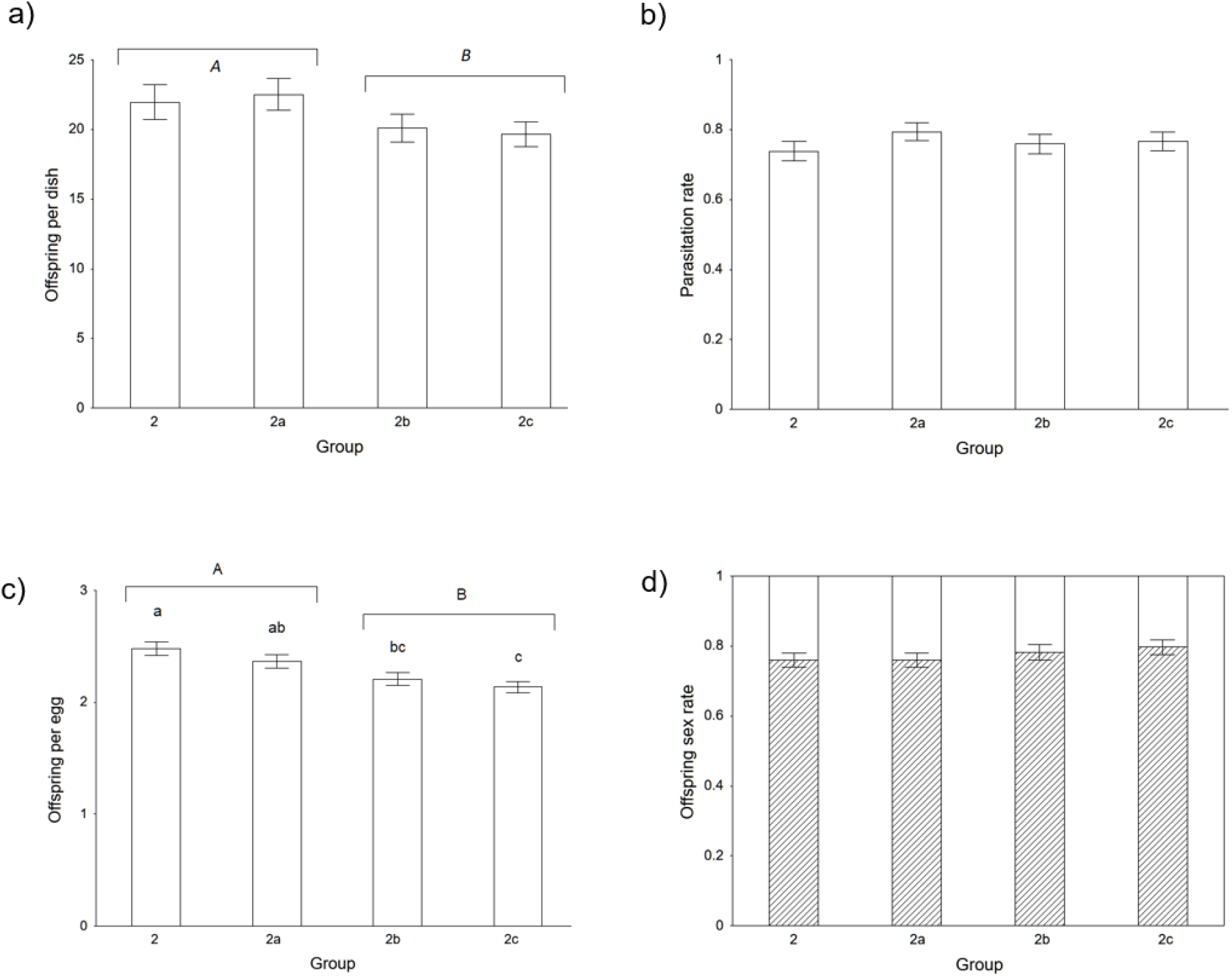
The relationship between the groups of wasps and the total number of offspring per dish (a), the total number of parasitized host eggs per dish (b), the offspring per one host egg (c) and the percentage of female offspring (d).

## DISCUSSION

### Effect of different numbers of founders

One of the indicators of superparasitism for gregarious parasitoids is a higher clutch size than under optimal laboratory conditions^e.g.40,41^. In this study, we reported this hypothesis for *A. flavipes*, since the clutch size increased with a higher number of founders. Such a situation generates stronger competition for food resources among the larvae^11,41^ and subsequently causes a future smaller offspring body size^11,12^ and lower fertility^13,8^. In our previous study Samková et al.^8^, we suggested that in the generally known trade-off scheme, the female decision between more small or fewer larger offspring does not apply to *A. flavipes* because large females have more offspring compared to smaller females and their body size is determined by the clutch size, thus with a sufficient number of hosts, females can lay a lower clutch size and their offspring will be larger with higher fertility^8^. This situation is predicted for many other species of gregarious parasitoids^e.g.42,30^. However, in superparasitism, a female should use a trade-off between the total number and the body size of the offspring due to limitations by the host. The offspring body size of *A. flavipes* proved to be a crucial factor because females did not use the host to the maximum (max. clutch size is 7 offspring per host^31^) but instead chose to produce fewer larger offspring at the cost of reduced own total fertility. Our result is the same as the general trends, namely the fertility of gregarious parasitoids does not increase linearly with a higher number of founders, and the fertility per each female decreases^25,this study^.

When evaluating the fitness of parasitoids, two different theoretical approaches have been proposed, first only from the view of the mother’s fertility and the second considers not just the mother’s view but also that of her offspring’s fertility^43^. Based on our results with superparasitism, we suggest, for gregarious parasitoids, it is important to consider the second approach (mothers and also offspring fertility). As shown in Fig. 3, with the same number of hosts, if the founders retain fixed fertility, their offspring will be extremely small with reducing future fertility^8^ (Fig. 3a) and at a disadvantage compared to large individuals^44^. However, when each of the founders reduces her fertility by half (Fig. 3b), their offspring will be larger with a higher own fertility in the F2 generation, almost identical to the fertility of smaller offspring^8^. We showed that with a scarcity of food for larval development during superparasitism, the females increase individual offspring fertility by reducing their own fertility.

**Fig. 3.**
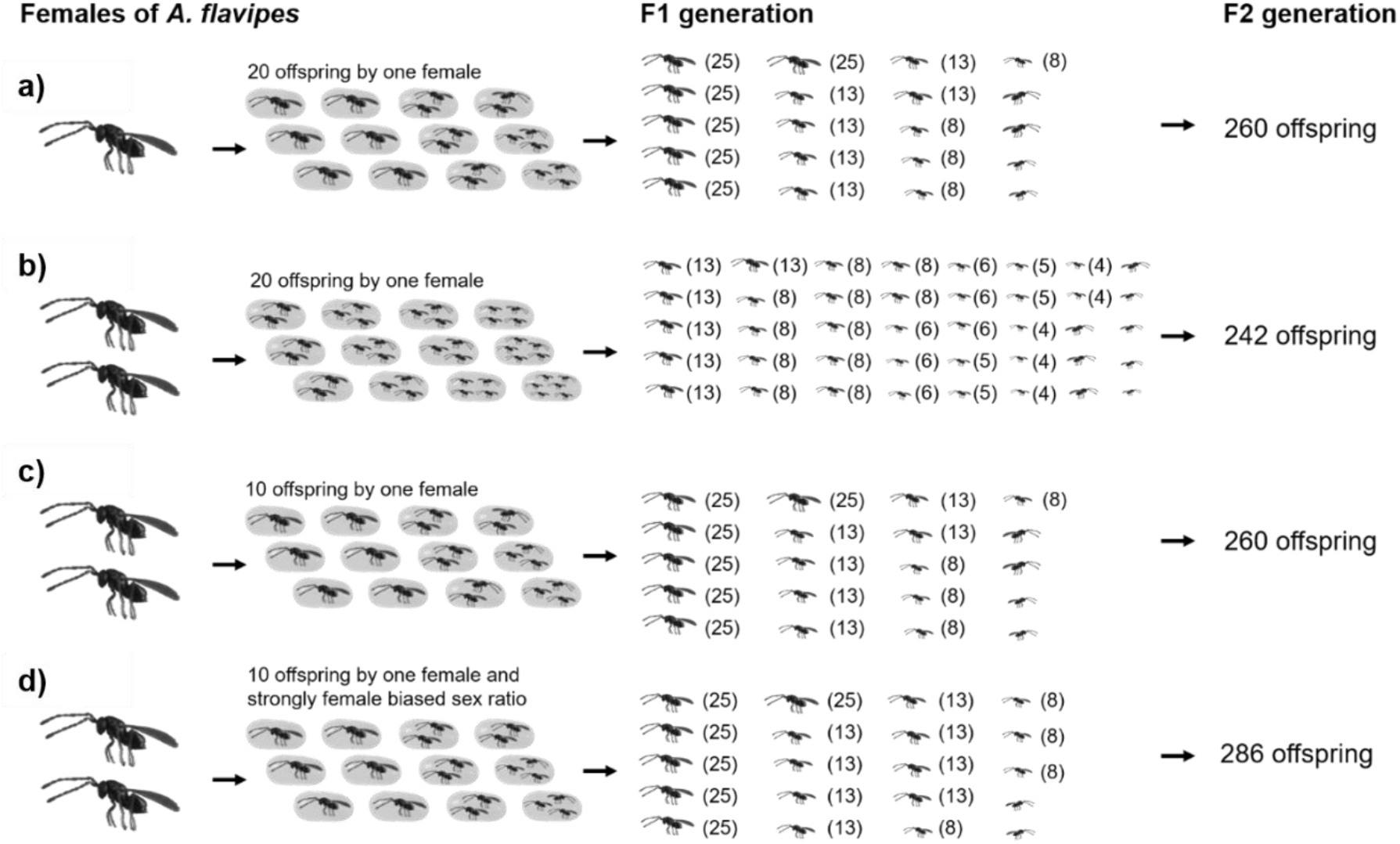
Based our previous results that the *A. flavipes* fertility increases continuously with the body size of the female and the body size depends on the clutch size^8^, we present a figure with the hypothetical fertility of the F1 generation. With a scarcity of hosts the females increase their individual offspring fertility by reducing their own fertility: (a) Under optimal conditions (12 host eggs to parasite, 20 offspring in the F1 generation), the mother female has hypothetical fertility 260 offspring in the F2 generation; (b) Two founders retain the same fertility (20 offspring per female), but due to the higher clutch size, the body size and fertility of their offspring decreases; (c) Each of two founders reduces her fertility by half (10 offspring per female) and due to reduced fertility their offspring will be larger with higher fertility; (d) Two founders reduce their fertility by half (10 offspring by one female) and the sex ratio of their offspring is strongly female-biased, thereby ensuring higher fertility of the F2 generation (d).

We are aware that our hypothetical fertility in the *A. flavipes* F2 generation presented in this study (Fig. 3) does not describe this phenomenon in detail. With large clutch size, fertility per each offspring decreases^8^, but the decrease in fertility can have different intensities in different species and may not be as simple as presented in Fig. 3. However, we showed that for gregarious parasitoids with a body size-fitness correlation such as A. *flavipes*^15^, it is necessary to assess its fitness from the view of both mothers and their offspring. In addition to higher fertility^11^, larger female offspring can also have larger eggs^4^, longevity^45^ and higher efficiency in the search for food resources or hosts^44,11^. Similarly, larger males have a longer life span^46^ and can defend their territory more successfully^47^ or be more successful in mating^11^. Due to reducing their fertility by half, the females retain half of their eggs for the possibility that they will find another host^25^ or for energy to extend their life span^48^ or to search for other hosts^49,50^.

Aside from fertility reduction in superparasitism, females with haplodiploidy sex determination^7^ can also change their offspring sex ratio^51,3^. Under normal conditions, the sex ratio of *A. flavipes* offspring is female biased 3:1 (females: males^31^) or 4:1 (females: males^Samková et al. unpubl.^), the same as in many other species of gregarious parasitoids^e.g.52,53,54^. In our previous study, we found the *A. flavipes* females change their reproductive strategy during a lower host population density, and in the presence of host egg predators, they increase their clutch size, but retain a stable female-biased offspring sex ratio^Samková et al. unpubl.^. We, therefore, hypothesised that with a higher clutch size and a higher degree of superparasitism, the females would retain a constant offspring sex ratio. Contrary to our expectations, with a higher degree of superparasitism, the proportion of female offspring increased, and the mothers reduced competition among their sons^55^ and obtained higher fertility in the F2 generation from their female offspring (the hypothetical model is shown in Fig. 4d). This result is consistent with the study of Montoya et al.^56^ on the species *Diachasmimorpha longicaudata* (Ashmead, 1905) and is in contrast to studies where the offspring sex ratio was not influenced^57,58^ or male-biased^59,60^. We propose that female-biased offspring is another reason why mothers do not lay the maximum number of offspring into one host, because it would severely reduce the offspring’s body size, and fertility in females is more affected by body size than in males^14^.

**Fig. 4.**
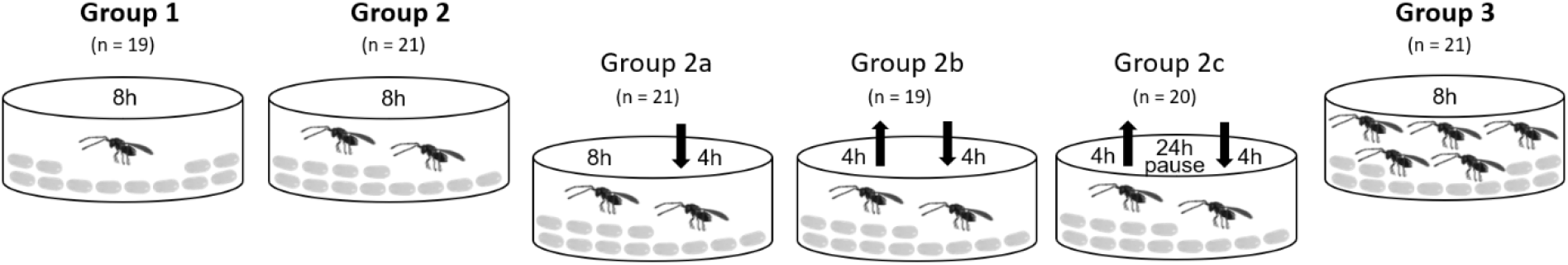
Six groups of *A. flavipes* wasps. *Group 1*: every female had for parasitation host eggs for 8 h; *Group 2:* two females had for parasitation host eggs for 8 h; *Group 2a*: The first female had for parasitation host eggs for 8 hr, after 4 h, a second female was added into the same Petri dish for 4 h; *Group 2b*: The first female had for parasitation host eggs, after 4 h, the first female was removed and a second female was added into same Petri dish for another 4 h; *Group 2c*: The first female had for parasitation host eggs, after 4 h, the first female was removed and after 24 h a second female was added for another 4 h; *Group 3*: five females had for parasitation host eggs for 8 h.

In this figure, we show the hypothetical fertility of the F1 generation, with half the amount of food (host) for larval development, the wasps’ body size and fertility are halved, but it must be emphasized that the decrease in fertility with decreasing body size, due to half the amount of food for larval development, will be different for different species (clutch size / fertility (rounded)) – 1/25; 2/13; 3/8; 4/6; 5/5; 6/4).

### Different parasitation timing and interactions

Although the event of parasitation by more founders can be recognized only by using genetic methods^58,61^, we assumed certain trends in the reproductive strategy of *A. flavipes* with a different design for parasitation. We expected possible differences in fitness characteristics for *Group 2b*, in which the female was alone with host eggs and had a short latency between parasitation compared to *Group 2c*, with a similar design, but the latency between parasitation was 24 h. The long latency caused the occurrence of *A. flavipes* larvae in the host eggs parasitized by the first founder before laying by the second founders because *A. flavipes* larvae emerge 19 hours after parasitation^62^. The larvae of the second female may be at a disadvantage compared to the larvae of the first female^14,63^ and the survival of a second clutch laid into an already parasitized host is generally low^34,39^, especially with a long latency between parasitation^41^. For example, the offspring of second females of *Trichogramma euproctidis* (Girault, 1911) were successful when the intervals between parasitation of the first and second founders were short (0–1 h), but if the intervals were 17–19 or 27–29 h the offspring of the second founders were outcompeted by the offspring of the first founders^21^. Our measured fitness characteristics did not statistically differ between *Group 2c* (longer latency) and *Group 2b* (short latency). We assumed successful superparasitism of *A. flavipes* due to their tolerant larvae without the effect of long latency between parasitation. However, in the future, other experiments with longer latency periods will be interesting, because generally, the longer the time interval between parasitation, the more the second clutch is disadvantaged^64,65^. For any species with a long time between parasitism, the second founder hyperparasites the larvae of the first founder, and the first larva consume all of the host tissues and occupy most of the host egg (found in the families Trichogrammatidae^21^, Mymaridae^66^ and Bethylidae^67^), and for *A. flavipes* this latency is 96 hours^62^.

When we compared the fitness characteristics of the four groups, the founders, which had visual information about the other female, had a larger clutch size than founders that were with the host eggs separately (see results). This finding is even more apparent when comparing the fitness characteristics of the emerging groups. The founders with information on the second females (*2 + 2a*) have a higher clutch size and higher fertility compared to the females left separately with the host eggs (*2b + 2c*). These results would indicate that the founders change their reproductive strategy according to information about other females^14^; however, we must be careful in interpreting these results, because there were different times between the groups that the wasps were with the host eggs. The offspring sex ratio and the total number of parasitized eggs for the four groups (*2, 2a, 2b, 2c*), as well as the emerging groups (*2a+2b* and *2b+2c*), were not different. The sex ratio of the *A. flavipes* offspring corresponds to the study by Shuker & West^58^, in which the sex ratio of *Nasonia vitripennis* (Walker, 1836) remained unchanged, irrespective of whether the superparasitizing founders laid eggs alone or while surrounded by conspecifics.

## CONCLUSION

Based on our results, we point out the fact that for determining the fitness of gregarious parasitoids with a body size fitness correlation, it is necessary to take into account fertility in both the F1 and F2 generations. Especially with a scarcity of hosts, the maternal female increases the fertility of the individual offspring by reducing her own fertility. In this study, we determined the hypothetical fertility of the F2 generation using a hypothetical model according to clutch size, because we know that with a larger clutch size, the body size of the offspring and their future fertility decreases^8^. However, in future work on this topic, we propose to model the exact F2 generation fertility in gregarious parasitoids with a body size fitness correlation using the actual body sizes of all offspring. We propose the offspring body size can be variable independently of clutch size with superparasitism or with a different lantency between two parasitations as has been shown for solitary parasitoids (different offspring body sizes with superparasitism^42,68^; with different latencies between parasitations^21^). Additionally, we confirmed the general trend that with a higher number of founders and a scarce number of hosts, the clutch size increases but the fertility per female decreases. We also found other trends in the changes of the reproductive strategy of *A. flavipes* in relation to a limited number of hosts with more founders. However, to clearly determine changes in reproductive strategies with different parasitation timings and interactions, we propose to use genetic methods to determine maternity in the future.

## MATERIALS AND METHODS

### Parasitic wasps

Parasitic wasps, *A. flavipes*, were collected from host eggs (*Oulema* spp.) from the end of April until the end of June 2019 in cereal fields (barley) in Prague-Suchdol, Czech Republic (GPS: 50.1367N, 14.3638E). The parasitized host eggs were stored in Petri dishes with moist filter paper until the adult wasps emerged. These “wild” wasps were used as an initial population for rearing the next generations of parasitoids in an environmental chamber with conditions of 22±2°C, a 40–60% relative humidity and 24-hour light. All “next generation” females and males of *A. flavipes* used for the experiments were at most 24 hours old (post-emergence). Mated females (not older than 24 hours post-emergence) were placed in Petri dishes with host eggs. Before the start of the experiments and during the experiment, all females were not fed, but they had free access to water (taken from Samkova et al.^8,15,69,70^).

### Host species

In our experiments, we used the *Oulema* species complex including two very ecologically close species *O. duftschmidi* Redtenbacher, 1874 and *O. melanopus* Linnaeus, 1758, because these two host species occur together in the same localities at the same time^71,70^ and are determinate only on the basis of genital preparations^72^. We consider it safe to use these two species in our experiments because their host eggs are not different in length or weight^8^, the wasps *A. flavipes* are not host-specific^70^ and wasp offspring from these two host species are not different in size and show no morphologic abnormalities^8,Samková et al. unpubl.^. The *Oulema* species complex was used in our previous studies^8,15,69,70^ and other studies^e.g.31,73,62,74^, because the *O. duftschmidi* species was only described and separated from *O. melanopus* in Bezděk & Baselga^72^. Here, the host culture was established from adults collected at the same time as host eggs in two localities in the Czech Republic (in Prague-Suchdol [identical to those of the parasitic wasps] and Police nad Metují [GPS: 50.5277N, 16.2456E]), using net sampling or individual sampling. The *Oulema* species were kept in plastic boxes (16.7 × 17.4 × 6.4 cm) with moistened filter paper. Adults were fed grain leaves and had unlimited access to water. The *Oulema* species laid their eggs on cereal leaves in an environmental chamber at 22±2°C, relative humidity of 40–60% and 16:8 L:D cycle. Host eggs were removed along with a 1 cm long piece of leaf and then used in the experiments. *A. flavipes* females refuse host eggs older than 72 hours^31^ so in our experiments, we used eggs not older than 24 hours (modified from Samkova et al.^8,15,69,70^).

#### Laboratory experiments

All laboratory experiments were performed in Petri dishes (diameter 8.5 cm) in a thermal cabinet at 22±2°C, relative humidity of 40–60%, and a 24 h light photoperiod. Individual host eggs were removed on the 9th or 10th day after parasitation and placed in 1.5 ml plastic tubes and stored at the same temperature as in the thermal cabinet (modified from Samkova et al.^8,15,69,70^).

#### Experimental design

Petri dishes (121 in total) were divided into six groups. Twelve host eggs and different numbers of founders were placed into each dish, and each female spent a different amount of time in the Petri dish (see below, Fig. 4).

##### Group 1

One female had for parasitation 12 host eggs for 8 h. This group simulates the control group, with a respectively lower population density of wasps.

##### Group 2

Two females had for parasitation 12 host eggs for 8 h. This group simulates a medium population density of wasps.

##### Group 2a

The first female had for parasitation 12 host eggs for 8 h; after 4 h, a second female was added into the same Petri dish for 4 h. The second female has visual information about the first female.

##### Group 2b

The first female had for parasitation 12 host eggs; after 4 h, the first female was removed, and a second female was added into the same Petri dish for another 4 h. The second female had no visual information about the first female.

##### Group 2c

The first female had for parasitation 12 host eggs; after 4 h, the first female was removed and after 24 h and a second female was added for another 4 h. This group represented the group with a long latency between parasitation when the second female has no information about first female.

##### Group 3

Five females had for parasitation 12 host eggs for 8 h. This group simulates a high population density of wasps.

### (1) Different number of founders

Using three different groups of wasps (*Groups 1, 2* and *3*), we measured the effect of a different number of founders on their reproductive strategy (n = 61). The total number of parasitized eggs and the total number and sex ratio of offspring and clutch size were recorded for every Petri dish as characteristics of the wasps’ reproductive strategy (Supplementary material 1).

### (2) Same number of founders with a different design

Using four groups of wasps (*Group 2, 2a, 2b, 2c*) and their combination (*2*+*2a* x *2b*+*2c*) we measured changes in reproduction strategy with relation to a different design of parasitation (n = 81). The total number of parasitized eggs and the total number and sex ratio of the offspring and clutch size were recorded for every Petri dish as characteristics of the wasps’ reproductive strategy (Supplementary material 1).

#### Statistical data processing

Software R version 3.6.1^75^ was used for all statistical analyses. Values for whole Petri dishes (parasitized egg rate, offspring per dish) were processed in linear models (LM, for dependent variables that were normally distributed or transformed to a normal distribution) or generalized linear models (GLM, for dependent variables in other than a normal distribution—specifically, in binomial, Poisson and Gamma distributions of our data). All of the models included group (see Experimental design) as a single fixed factor.

When the analyses involved values of individual eggs (offspring per parasitized egg, offspring sex rate), mixed linear / generalized linear models (LMM/GLMM, package lme4^76^) were used, including group as a fixed factor and the individual label of each Petri dish as a random factor. Significance of the effect of the “group“ variable was assessed by comparison of the original model with a null model, which included only the random factor as an explanatory variable, by ANOVA.

## ACKNOWLEDGEMENTS

We would like to thank the colleagues who provided us with technical equipment: Jana Mazáková, Marie Maňasová, Jana Wenzlová and Miloslav Zouhar. This work has been supported by the Ministry of Agriculture of the Czech Republic, institutional support MZe-RO0418 and from programme NAZV No. QK1910281 (MZe ČR) (both to JS).

